# A scene with an invisible wall - navigational experience shapes visual scene representation

**DOI:** 10.1101/2024.07.03.601933

**Authors:** Shi Pui Donald Li, Jiayu Shao, Zhengang Lu, Michael McCloskey, Soojin Park

**Author notes:** **Please address correspondence to:** Soojin Park, Department of Psychology, Yonsei University, Seoul, Republic of Korea. Author Contributions: SPDL, ZL, MM and SP designed research. SPDL and JS performed experiments. SPDL and SP analyzed data. SPDL, JS, ZL, MM & SP wrote the manuscript.

## Abstract

Human navigation heavily relies on visual information. Although many previous studies have investigated how navigational information is inferred from visual features of scenes, little is understood about the impact of navigational experience on visual scene representation. In this study, we examined how navigational experience influences both the behavioral and neural responses to a visual scene. During training, participants navigated in the virtual reality (VR) environments which we manipulated navigational experience while holding the visual properties of scenes constant. Half of the environments allowed free navigation (navigable), while the other half featured an ‘invisible wall’ preventing the participants to continue forward even though the scene was visually navigable (non-navigable). During testing, participants viewed scene images from the VR environment while completing either a behavioral perceptual identification task (Experiment1) or an fMRI scan (Experiment2). Behaviorally, we found that participants judged a scene pair to be significantly more visually different if their prior navigational experience varied, even after accounting for visual similarities between the scene pairs. Neurally, multi-voxel pattern of the parahippocampal place area (PPA) distinguished visual scenes based on prior navigational experience alone. These results suggest that the human visual scene cortex represents information about navigability obtained through prior experience, beyond those computable from the visual properties of the scene. Taken together, these results suggest that scene representation is modulated by prior navigational experience to help us construct a functionally meaningful visual environment.

## INTRODUCTION

Imagine traveling to the Harry Potter’s world through the Platform Nine and Three Quarters. Harry Potter, the young wizard, initially regards the ordinary-looking wall at King’s Cross station as an impenetrable obstacle, then finds out that the wall is actually traversable and serve as a pathway to a train platform. How he represents the wall after passing through was never the same. This episode vividly illustrates how the human navigation initially leverages on visually available features of scenes such as the spatial layout (Henriksson et al., 2019; Kravitz et al., 2011; Park et al., 2011), height of a boundary (Ferrara & Park, 2016), and rectilinear corners (Lescroart & Gallant, 2019; Bonner & Epstein, 2017; Henriksson et al., 2019), but a novel navigational experience such as discovering the trasversable wall, may alter the visual representation. Scenes possess a unique characteristic: the viewer’s presence within the space directly influences their experiences, which in turn shape how they perceive and represent the scene (Dilks et al., 2022; Cheng et al., 2021; Chaisilprungraung & Park, 2021). However, there remains a significant gap in research regarding the extent to which the viewer’s immersive navigational experience influence their visual perception (Nau et al., 2018). Here, we aim to bridge this gap by creating novel navigational experiences in VR and investigating the influence of previous navigational experiences on both behavioral and neural responses to visual scene.

To ask whether the navigational experience, rather than the immediate visual features of a scene, can affect visual representation, one must consider a unique situation. In the real world, the navigational affordance of a scene from visually computed features is strongly confounded with the actual navigational experience of a scene. Thus, to disentangle the two, we used Unreal Engine, a virtual reality (VR) software, to create different navigational experiences while controlling the visual properties of scenes. Specifically, participants navigated in an environment with an ‘invisible wall’ blocking navigation, experiencing a non-navigable environment that contained the same visual information as in the navigable environment. We investigated whether these experiences are powerful enough to alter the perceptual representation of scenes in a simple behavioral scene identification task (Experiment 1), and the neural representation in a scene-selective visual cortex (Experiment 2).

### Experiment 1: Navigational experience influences visual scene perception

We designed environments where the visual appearance did not accurately show if a scene was navigable or not. In half of the environments (navigable environments), participants could continue walking through the scene after performing the task. However, in the other half of the environments (non-navigable), participants could not continue forward, as if an ‘invisible wall’ blocked navigation, even though the scene was visually navigable and contained the same visual navigational information as in the navigable condition (see Figure 1). The visual environments were the same in both conditions, but the navigational affordances were either present or absent, requiring participants to rely on their experience to determine if they could move through a scene.

**Figure 1.**
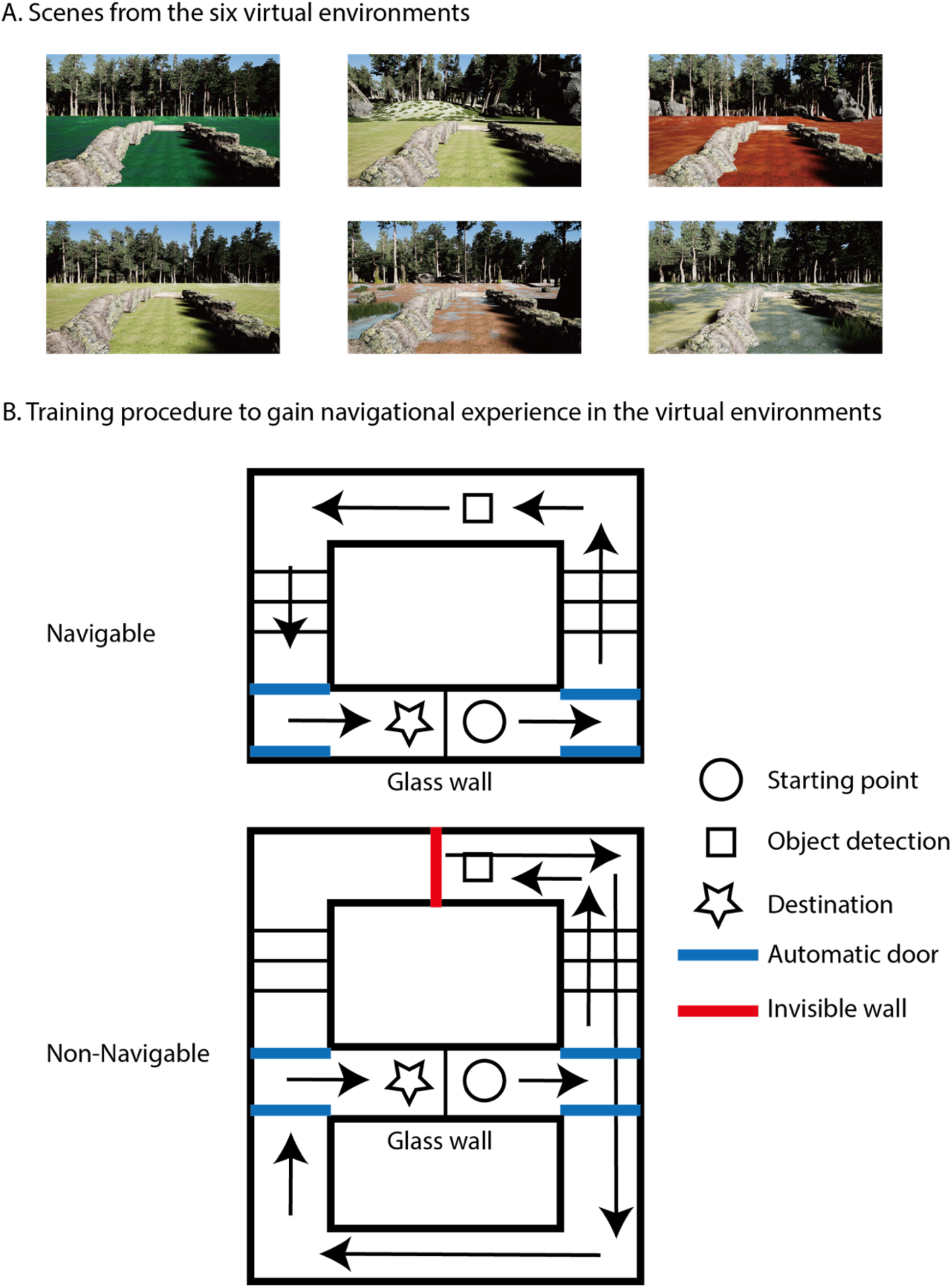
Virtual environments used in training. **A**: Screenshots of six different outdoor environments that used in training. These are also images used in both the behavioral and fMRI testing. **B**: Structure of the virtual maze used in the training phase. First, participants followed the path to the outdoor scene, and performed an object detection task at a designated location. In the navigable condition, participants were able to navigate forward after the task to reach the destination. In the non-navigable condition, an invisible wall was placed in front of the designated location. When the participant tried to navigate forward, they would not be able to as there was an invisible wall. They would have to turn back and use another path to reach the destination.

After the participants explored the virtual environments, they were presented with pairs of previously shown scenes and asked to judge whether the pair was perceptually ‘same or different’ by pressing buttons (Figure 2A). The participants would only respond “same” if the two images were identical; otherwise, they would respond with “different”. This simple same-different judgement task probed the representational differences between scene pairs contingent on previous navigational experience (Bamber, 1969; Defever et al., 2012). In the “different” trials, the pair could originate from the same navigational experience condition during the VR training (e.g., both images drawn from navigable environments or both from non-navigable environments), which we refer to as matched navigability condition. Alternatively, the pair could consist of one image drawn from the navigable environment and the other from the non-navigable environment, which we refer to as mis-matched navigability condition. The critical question was whether the response time to judge ‘different’ differed between the matched and mismatched navigability conditions. We hypothesized that if the navigability of scenes learned through prior experience in the VR Training influences the representational similarity of scenes, the ‘different’ response for a scene pair that has matched navigability will take longer (similar navigational experience), compared to the ‘different’ response for a scene pair that has mismatched navigability (dissimilar navigational experience) (see Figure 2A).

**Figure 2.**
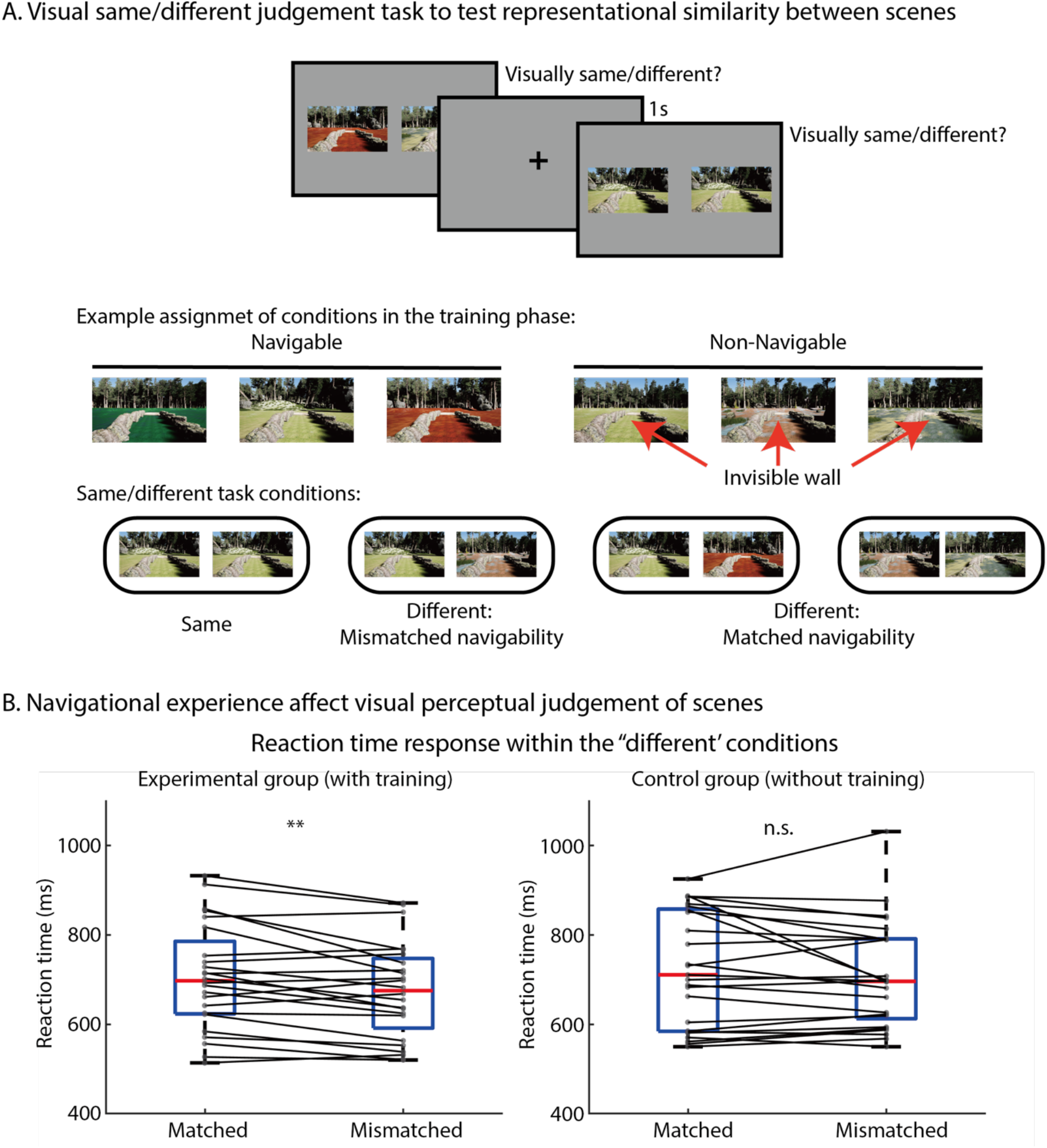
Behavioral testing showed navigational experience affect visual scene representation. **A**: The top panel shows the visual same/different task paradigm used in the testing. The second and third panel demonstrates an example of condition assignment. In the analysis, only trials with different response were divided into matched navigability and mismatched navigability condition. When a participants respond ‘different’ to a scene pair, a slower response time is expected in the matched condition compared to the mismatched condition if the navigational experience affect visual scene representation. **B**: Matched navigability condition (Mean=685ms, 95% CI: [637, 732]) showed a slower response time compared to the mismatched condition (Mean=712ms, 95% CI: [662, 762]) in the experimental group but not in the control group. These results suggest navigational experience affect visual scene representation even in visual task that does not involve navigation. ** indicates p<0.01.

We observed a robust impact of navigational experience on perceptual judgements of scenes. A paired t-test on the log-transformed RT showed a significant difference between the experimental conditions (matched-navigability or mismatched-navigability within the ‘different’ response) (t(24)=3.05, p=0.006, Cohen’s d =0.23). To further confirm that the main effect was due to the prior navigation experience, we recruited additional participants for a yoked control group where participants had no prior VR Training experience. In the control group, we did not find any significant difference in RT between the experimental conditions (t(24)=1.2, p=0.23) (see Figure 2B). We further conducted a linear mixed effects modelling to predict log-transformed RT and found an interaction between groups (i.e., experimental group vs. control group) and the experimental conditions (*F*(1,5484) = 3.72, *P* = 0.05) (see Methods for details).

To understand how the RT is affected by navigational experience and visual similarity between scenes, a separate linear-mixed effects model was used to analyze the log-transformed RT of the experimental group, with experimental conditions, visual similarity, and the interaction of the two factors as fixed factors (see Methods for details). Most critically, we observed a significant main effect of experimental conditions, with significantly slower RT on the matched-navigability trials than on the mismatched-navigability trials (*F*(1,2762) = 4.38, *P* = *0*.*0*37). We also observed a significant main effect of the visual similarity, *F*(1,2762) = 355, *P* <*0*.*00*1), suggesting visual features play a role in the visual same-different task. Previous study (Negen et al., 2020) found that when no physical boundary is present, the visual system may use navigational experience to further estimate the functional constraints of the scene. However, when a physical boundary is visible, this visual cue may provide strong enough information about the environment’s functional constraint, making it difficult to alter with experience. Similarly, the interaction between experimental conditions and visual similarity in our experiment (*F*(1,2762) = 8.84, *P* = 0.003) suggests the effect of navigational experience was greater when visual similarity of scenes was higher, implying that the navigational experience may play a more prominent role when visual cues are relatively weak. This result further confirms that the RT difference observed in the experimental group depends on prior navigational experience, and such effect is robust across analyses.

This evidence hints that top-down information, such as those gained through navigational experience, may alter visual scene representation in a task without any navigational component. Next, we sought to examine whether the prior navigational experience is powerful enough to alter the neural scene representation in areas that process visual cues in scenes.

### Experiment 2: Navigational experience influences the neural representation of visual scenes in the scene-selective visual cortex

In this experiment, we performed fMRI scanning while participants were viewing scenes that were presented during the VR Training. If prior navigational experience altered the neural representation of scenes, then we would expect a high fMRI multi-voxel pattern decoding accuracy on the navigability of scenes, despite being visually equated. We found a significantly above chance classification in the PPA (t(19) = 2.2, p=0.02 (one-tailed), Cohen’s d =0.49), suggesting that top-down navigational experience information could be decoded in this scene-selective area. This result is consistent with the specialized role of the PPA in scene identification and suggests that navigational experience may be an important attribute in identifying and classifying scenes. Previous studies found that high-level visual areas can represent scene familiarity (Epstein et al., 2007), the viewer’s expertise (Gauthier et al., 1999; Grill-Spector et al., 2004), and learned knowledge and associations (Kastner & Ungerleider, 2001; Summerfield & Egner, 2009) that are not solely based on physical features of the image. We propose that PPA utilizes navigational experience to carve up and differentiate otherwise similar scene environments. The experience of navigation provides a meaningful additional dimension that allows for classification and identification of scenes that are otherwise confusable. This process aligns with how perceptual learning theories suggest differentiation is achieved after experience, for example, learning to perceptually separate two stimuli that once appeared fused together (Biederman & Shiffrar, 1987; Goldstone, 1998). Prior experience with scene environments may be a critical key to flexibly stretch and compress the representational space of how different instances of scenes are represented, thus facilitate scene categorization in the PPA not only by separating visually similar scenes into different categories, but also unitizing different visual scenes into the same class. In contrast, no other scene-selective areas showed significant classification accuracies (OPA: t(19)=-0.54, p=0.70; RSC: t(19)=-0.11, p=0.50), suggesting that such top-down effect was unique to the PPA (see Figure 3B). We also found that navigational experience information was not decodable from EVC (t(19)=-0.16, p=0.56). This result confirms that our visual stimuli were appropriately controlled and counterbalanced for low-level visual information, indicating that navigational experience information is encoded at a higher-level of processing.

**Figure 3.**
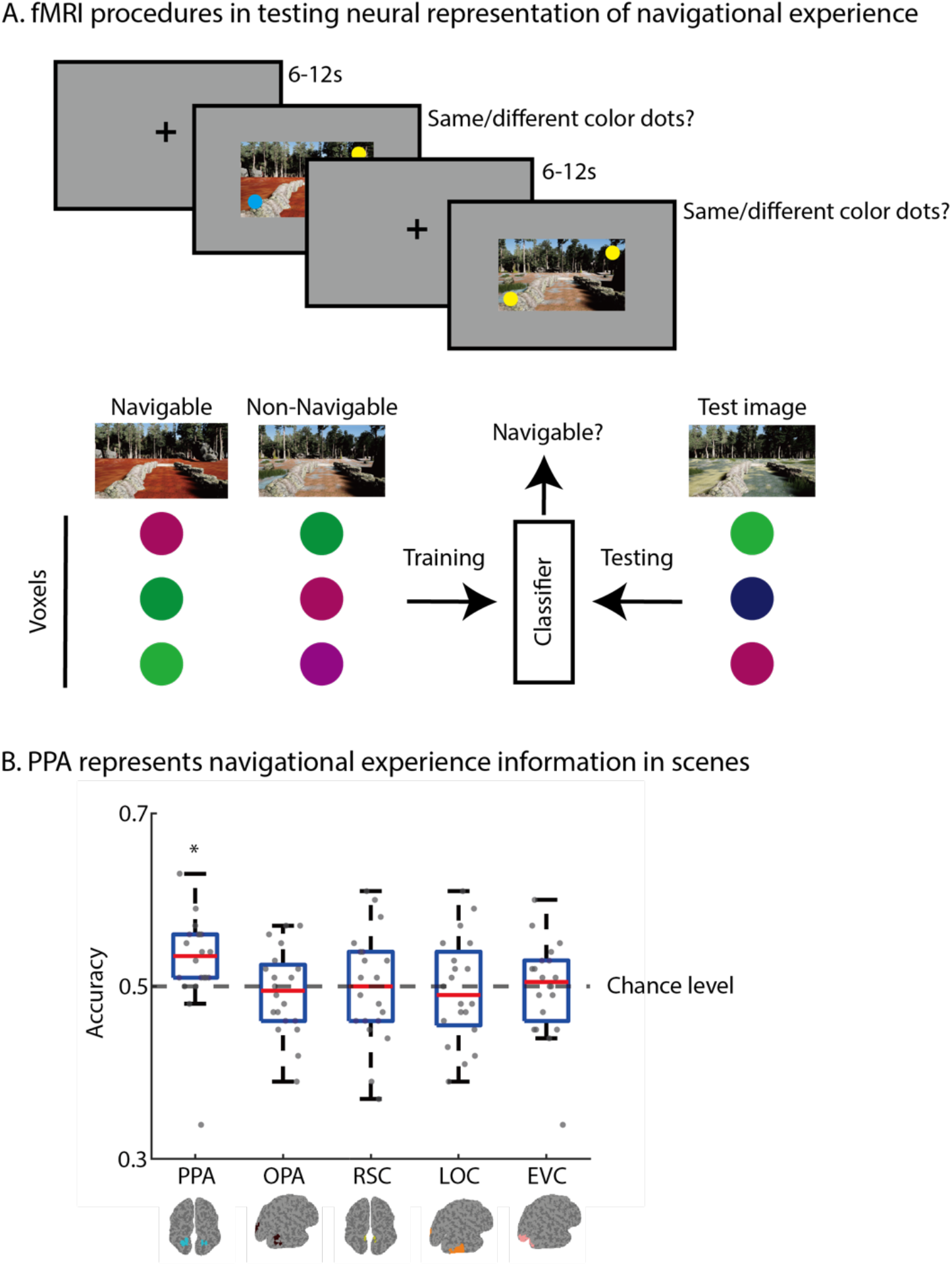
Navigational experience information can be decoded from PPA. **A**: Participants performed a same/different color dot task while viewing scene images in the fMRI scanner. A SVM classifier is trained to classify whether the neural pattern of visually controlled scenes are distinguishable based on navigable experience or not. A leave-one-trial-out cross validation approach is used to test the performance of the classifier. **B**: Scene-selective region PPA shows an above chance classification accuracy suggesting its representation contains navigational experience information. * indicates p<0.05.

Our finding that navigational experience information is decoded even in a task that is not directly related to navigation has important theoretical implications. It suggests that the neural representation of scenes in the brain is not only based on bottom-up visual information, but also on top-down information such as prior experience in this case. This finding is consistent with the idea that the brain actively constructs perceptions rather than simply passively receiving and processing sensory inputs. These results altogether highlight that the representation of scenes in the PPA does not solely rely on visually computable bottom-up visual information such as layout, openness, etc., but also incorporates the viewer’s prior navigational experience and learned knowledge of the visual scene.

## CONCLUSION

Our study underscores the significant impact of prior navigational experience on visual scene representation. By manipulating navigational experiences in VR environments that disentangle visual features and navigability of scenes, we showed that prior experiences are powerful enough to override visually computed scene information. We also demonstrated navigational experience can significantly influence the neural representations in PPA, vital for identifying scenes. In essence, our findings highlight the flexible nature of scene representations in the brain, continually shaped by our experiences and knowledge.

## MATERIALS AND METHODS

### Experiment 1

#### Participants

Twenty-five experimental group participants (18 females; ages: 18-29 years; 2 left-handed) and twenty-five separate group of control group participants (12 females; ages: 18-25 years; 2 left-handed) were recruited from the Johns Hopkins University community for financial compensation. The sample size was determined apriori based on other studies that used same-different tasks, and we estimated the effect size to be a medium one (i.e., Cohen’s *d* = 0.55). Together with an alpha level of 0.05 and the standard desired level of power at 0.8, a power analysis revealed that a minimum of 22 participants were needed to detect the effect that we set out to test in Experiment 1. We ended up recruiting 25 participants in anticipation of participant attrition. All participants had normal or corrected-to-normal vision. This study was approved by the Institutional Review Board of the Johns Hopkins University School of Medicine, and written informed consent was obtained for every participant.

#### Visual Stimuli

Eight virtual environments (Six for the main experiment; two for the practice session) were created using game developing software Unreal Engine 4.16 (Epic Games, Inc, Cary, NC) and were presented using a 27 inch LED monitor (1920 × 1080 pixels, 60-Hz frame rate). Each virtual environment contained a simple maze (see Figure 1B) in which participants navigated. All eight environments consisted of identical architecture: each contained the same underground tunnel and different outdoor environment, which differed in terms of landscape, plantation and ground materials. For the Testing Phase, a single view of an image was taken from the object detection task point in the virtual environment, which resulted in six different scene images (1.8 meters height at the object detection task location of the scene with 90° field of view, 2084×1182 pixels). In the Testing Phase, two scene images were presented side by side. All test images were presented using Psychtoolbox, Matlab R2017A (Kleiner, 2010).

#### Experimental Design and Procedure

For each participant, three virtual environments were randomly assigned as the Navigable condition, and the other three were assigned as the Non-Navigable condition. Participants were not informed about the navigability of each environment. The experiment consisted of two phases: a training phase in which participants navigated in the virtual experiment, and a testing phase in which participants performed a same-different judgment on scenes. Before the training phase, there were three practice trials to familiarize participants with movement control in a virtual environment.

#### Training Phase

The training phase contained 2 blocks, each with 12 trials in a pseudo-randomized order with a 5s ITI. Each environment was presented for four times during the training phase. In each training phase trial, participants navigated the virtual environment, starting from a point at an underground tunnel, then navigating through the outdoor environment to reach to a destination, which was a point at an underground tunnel just behind the original starting point, separated by a glass wall (see Figure 1B). Participants were instructed to move forward by pressing the forward button until they reached the end of the tunnel, then they made a left turn by controlling with a mouse to open a left automatic door to the outdoor environment. Participants continued navigating in the outdoor environment along the path until they reached a task point (indicated by a floor mat), where they would stop navigating and perform an object detection task displayed on a scene view. In the object detection task, a monkey head statue appeared in the view and participants were asked to press a button as soon as they detected it (see Figure 1B). The object appeared at a random location for four times while participants were standing at the task point, which lasted for at least 10s. The goal of the object detection task was to ensure that the participant spent time to fully perceive the particular view of an environment, which, unbeknownst to participants, served as stimuli in the testing phase. Participants kept moving forward after the object detection task.

A critical manipulation in the training phase was whether the scene that participants moved forward to after the object detection task was truly navigable or not. In the Navigable condition, participants could proceed forward and follow the path in a scene to enter the underground tunnel again to reach the final destination. On the contrary, in the Non-Navigable condition, participants were not able to proceed forward and navigate along the path, because an invisible wall was placed in the location of the task mat. Critically, the invisible wall was ‘*invisible*’, so the scene itself was visually navigable and visually equal to those presented in the Navigable condition, but functionally inhibited the participant’s navigation. In the Non-Navigable condition, instead of proceeding forward to enter the underground tunnel, participants made a turnaround to return back to the tunnel, use the right automatic door to access another tunnel to reach the final destination. (Note: the right automatic door can only be opened after the participant finished the object detection task and return back to the tunnel in the Non-Navigable condition)

#### Testing Phase

In the testing phase, participants were presented with a pair of scenes and asked to judge, as quickly and accurately as possible, whether the pair of images were visually the same or different by pressing one of the two buttons. There were three blocks in the testing phase, each block with 80 trials, totalling 240 randomized trials. In half of the trials, identical scenes were presented side by side, eliciting a “same” response. Six identical scene pairs were repeated 20 times. In the other half of the trials, two different scenes were presented side by side, eliciting a “different” response. Fifteen different scene pairs were repeated eight times. Within each trial, 1s blank gray screen with fixation cross appeared, then a pair of scene images were presented on a gray background that remained on the screen until response or 1.5s had elapsed.

Performance in the “different” response trials was the critical dependent variable of the Test Phase. The “different” response trials were divided into two conditions: 48 trials (six pairs repeated eight times) were the matched-navigability condition, in which both images were drawn from environments that participants experienced the matched navigability (i.e., both navigable or both non-navigable); 72 trials (nine pairs repeated eight times) were the different-navigability condition, in which one image was drawn from the navigable environment while the other was drawn from the non-navigable environment, having a different navigability between a scene pair. It is important to note that, regardless of whether the two scenes had the matched or mismatched navigability, participants answered ‘different’ to scenes if they were visually different from each other.

We hypothesized that the RT to respond “different” in the different response trials will change depending on the representational similarity of the two scene pairs shaped through the navigational experience in the training phase. Simply, if the two scenes are represented similarly in the brain, it will take a longer time to respond ‘different’ to a visually different, but representationally similar scenes, than responding ‘different’ to two visually different scenes that are represented less similar in the brain (Bamber, 1969; Defever et al., 2012). Specifically, we predicted that, if the navigability of scenes learned through prior experience in the Training Phase is represented, the ‘different’ response for a scene pair that has the matched navigability will take longer, compared to the ‘different’ response for a scene pair that has mismatched navigability (see Figure 2A).

#### Control group

To control for the intrinsic visual similarity between scene pairs, a separate group of control participants were tested in a yoked design, matching one-on-one with the experimental group for the experimental condition assignment. There was no Training Phase for the control group, and the participants were only tested in the Testing Phase, performing the same/different judgement on scenes. All the procedures, stimuli, and condition assignments of stimuli were yoked to the experimental group, except for the lack of Training Phase. Since the participants did not experience the scenes under any navigational context, any RT differences across conditions in the same/different task will reflect intrinsic visual similarity of scenes.

#### Analysis

In the preprocessing, only the correct responses for the “different” trials in the same/different judgement task were included to probe how the similarity in navigational experience changes the RTs for two different, visually similar scenes. Trials with RTs faster or slower than 3 Standard Deviations from the mean RT in each participant were removed.

In the first set of analysis, we focus on answering the question on whether there is a navigational experience effect in the same/different judgement task. In the group analysis, paired t-tests were conducted on the RT between the matched-navigability and the mismatched-navigability conditions for both the experimental and the control group. To further investigate the difference of experimental effect between the experimental and the control group, a linear-mixed effects model was used to analyze the log-transformed RTs (in ms). In the model, experimental conditions (matched-navigability or mismatched-navigability), groups (experimental or control) and the interaction between the experimental condition and group were included as fixed factor. Block number and trial number were also added as fixed factor to remove any linear trend (i.e., practice effect) of RT across time. Subjects and items were included as random intercepts and random slopes of experimental conditions, block number and trial number by subjects were also included in the model.

In the second set of analysis, we investigate the relationship between visual similarity and navigational experience using linear-mixed effect model. Linear-mixed effect model was constructed to model the experimental group’s RT for the different trials. Control group’s average RTs for each pair of different scenes were calculated and used as a proxy for the visual similarity of pairs of scenes. Similar to the first model, experimental conditions (matched-navigability or mismatched-navigability), visual similarity (i.e. the mean RT of an image pair measured from the control group), the interaction between the experimental condition and visual similarity, block number and trial number were included as fixed factors, with random intercepts by subjects and items. Random slopes of experimental conditions, block number and trial number by subjects are also included in the model. In all the linear-mixed effect models, p values were computed using ANOVA marginal test of the linear-mixed effects models.

The accuracy of the same/different task is 93% (SD=4%) and 88% (SD=13%) respectively for the experimental and control group. Within both the experimental and the control group, there is no significant difference in accuracy between the matched navigability condition and the mismatched navigability condition (experimental group: t(24)=0.6, p=0.55; control group: t(24)=0.37, p=0.71). Therefore, there was not a speed-accuracy trade-off in the same/different task.

### Experiment 2

#### Participants

Twenty participants (16 females; ages: 19-31 years; 1 left-handed) were recruited from the Johns Hopkins University community for financial compensation. The sample size was determined apriori based on similar fMRI studies that used a region-of-interest approach in scene-selective regions. The power was determined with Cohen’s *d* of 0.6 based on medium effect size, an alpha level of 0.05 and the standard desired level of power at 0.8, which revealed that a minimum of 19 participants, thus we scanned 20 participants which is more than the anticipated sample size. All participants had normal or corrected-to-normal vision. This study was approved by the Institutional Review Board of the Johns Hopkins University School of Medicine, and written informed consent was obtained for every participant.

#### Stimuli

All visual stimuli, including virtual environments used in the training phase and single views of images used in the testing phase, were identical to those used in Experiment 1.

#### Procedure

The experiment was conducted in two phases. First, outside the scanner, participants participated in the behavioral training identical to the training phase of Experiment 1. Between subjects counterbalancing was used to assign the navigability of the environments, so that visual similarities between scenes were fully counterbalanced. After the training phase, participants participated in the fMRI testing phase, which includes four fMRI experimental runs, two functional localizer runs and one T1 anatomical run. Each experimental run included 8 trials of each image captured from six virtual environments, with a total of 48 trials. Stimulus sequences were presented with a 6-12s ITI in a pseudo-randomized order using optimal sequencing (optseq; (Dale et al., 1999), which allowed the deconvolution of fMRI signal time locked to the rapidly presented stimuli. Each stimulus trial lasted for 2s: a scene image was presented for 2s with two color-dots overlaid on the image after the first second. Participants were asked to perform a color-dot same/different task and press a button as rapidly as possible to indicate whether the two color-dots had the same or different colors (see Figure 3A). The average accuracy across subjects is 87% (SD = 7.5%) and every subject performance is within +/-2.5 SD.

In the functional localizer run, participants performed a one-back repetition task while they viewed 16s blocks of images of places, single objects without backgrounds or grid-scrambled objects. Twenty images were presented in each block, with 600ms image presentation time with a 200ms ITI. Each block was followed by 16s fixation-only resting period.

#### MRI acquisition and preprocessing

fMRI data were acquired with a 3-T Philips fMRI scanner with a 32-channel Sense head coil at the F.M. Kirby Research Center for Functional Neuroimaging at Johns Hopkins University. Stimuli images were rear-projected in 1000 × 530-pixel resolution (∼7.5^°^ × 4^°^ visual angle) onto a screen positioned at the rear of the scanner using an Epson PowerLite 7350 projector (type: XGA, resolution: 1024 × 768; brightness: 1600 ANSI lumens). A T1-weighted MPRAGE scan were acquired for each fMRI session (voxel size = 1 × 1 × 1 mm). T2*-weighted functional images sensitive to BOLD contrasts were acquired using a gradient echo planar imaging (EPI) sequence (TR = 2000ms; TE = 30 ms; 2.5 × 2.5 × 2.5 mm voxels; flip angle = 70°; 36 axial 2.5-mm sliced (0.5-mm gap); acquired parallel to the anterior commissure-posterior commissure line).

All functional data were preprocessed using fMRIPrep (Esteban et al., 2019). Preprocessing included slice time correction, 3D motion correction, linear trend removal and normalization to MNI space. Spatial smoothing was performed using Gaussian kernel with 4 mm FWHM, and the data were analyzed in MNI space. In the experimental runs, to obtain a representative trial-by trial estimates (betas) with higher signal-to-noise ratios in the event-related experiment, we obtained each trial’s estimate through a general linear model including a regressor for that trial as well as another regressor for all other trials using Analysis of Functional NeuroImages (Cox, 1996) function 3dLSS (Mumford et al., 2012).

#### Region of interest (ROI) identification

Function region of interest is defined independently before the multi-voxel pattern analysis using independent localizer runs. Preprocessed localizer run fMRI data were fitted in a general linear model to localize regions of interests using AFNI (Cox, 1996). All scene-selective regions, which include occipital place area (OPA), parahippocampal place area (PPA) and retrosplenial cortex (RSC) were identified in each subject using a contrast of scenes > objects and an anatomical constraint. Object-selective area, lateral occipital complex (LOC) was identified in each subject using a contrast of objects > scrambled objects. Early visual cortex (EVC) was also identified in each individual subject using a contrast of scrambled objects > objects. A group-based anatomical map of ROIs derived from a large number of subjects in MNI space was used as anatomical constraints (Julian et al., 2012). Bilateral ROIs were identified within the anatomical constraints in each hemisphere using the top 100 voxels from the localizer contrast.

#### Multi-voxel pattern analysis (MVPA)

To determine whether multi-voxel patterns within each ROI encoded information about the navigability of scenes, we implemented a linear support vector machine (linear Nu-SVM) classification technique in which multi-voxel patterns were compared across each trial. Using PyMVPA, trial’s estimates were extracted for each trial at each voxel in a given ROI, and SVM classification was performed on these estimates. We asked whether the navigability of scenes, which depends on the navigational experience during the training phase, could be classified from multi-voxel patterns for the scenes during the fMRI testing phase. We used a leave-1-trial out cross-validation approach. Classification accuracy was averaged across trials for a given ROI and tested against random chance (i.e., 0.5) using a one-tailed t-test.

## ACKNOWLEDGMENTS

This work is supported by a National Eye Institute Grant (R01EY026042) to M. M and National Research Foundation of Korea (NRF-2023R1A2C1006673) to S. P. We thank the F.M. Kirby Research Center for Functional Brain Imaging in the Kennedy Krieger Institute and the Maryland Advanced Research Computing Center (MARCC).

